# GPI-anchored SKU5/SKS are maternally required for integument development in Arabidopsis

**DOI:** 10.1101/813733

**Authors:** Ke Zhou

**Affiliations:** FAFU-UCR Joint Center for Horticultural Biology and Metabolomics, Haixia Institute of Science and Technology, Fujian Agriculture and Forestry University, Fuzhou, 350002, China; Institut des Sciences du Végétal, UPR2355, CNRS, Saclay Plant Sciences, 91198 Gif sur Yvette Cedex, France

**Keywords:** Glycosylphosphatidylinositol, GPI-anchored protein, GPI-AP, Integument Development, Seed Morphogenesis, *Arabidopsis*

## Abstract

Glycosylphosphatidylinositol-anchored proteins (GPI-APs) play crucial roles in various processes in eukaryotes. In *Arabidopsis, SKS1, SKS2, SKS3* and *SKU5* from *SKU5/SKS* gene family could encode GPI-anchored proteins, and they were recently reported to regulate cell polar expansion and cell wall synthesis redundantly in roots. Here, we report that, they are also redundantly crucial for seed production and seed morphogenesis in *Arabidopsis* through regulating maternal integument development. Their *loss-of-functions* resulted in disrupted development of integuments that failed to protect embryo sacs from exposure to external space due to physical restriction, and lead to female gametophytic abortion. Interestingly, those less defective ovules could be fertilized and develop into seeds normally, however, their seed morphogenesis was largely affected.

Our research made SKS1, SKS2, SKS3 and SKU5 be not only the first class of GPI-anchored proteins that could regulate maternal integument development, but also the first class of proteins that could regulate cell polar expansion in both root and integument cells besides several MAPK cascade components. Our study also underlined the regulation by integument development in reproductive processes.

## INTRODUCTION

In eukaryotes, various protein precursors could be modified during maturation in endoplasmic reticulum and Golgi bodies, where their special hydrophobic carboxyl-terminal signal polypeptides could be cleaved and catalyzed to be replaced by a common glycosylphosphatidylinositol (GPI) oligosaccharide structure. It covalently links the lipid bilayer and the carboxyl-terminus of glycosylphosphatidylinositol-anchored proteins (GPI-APs), which allows these proteins to be attached on outer surface of plasma membrane (Kinoshita and Fujita, 2016; Oxley and Bacic, 1999; Stevens, 1995; Zhou, 2019b). Disruption of GPI moieties biosynthesis resulted in broad and severe defections during various developmental processes (Bundy et al., 2016; Gillmor et al., 2005; Kawagoe et al., 1996; Kinoshita, 2014).

In *Arabidopsis*, more than 300 GPI-APs from diverse families have been identified or predicted by bioinformatics or proteomic tools (Borner et al., 2003; Takahashi et al., 2016), among which, only a small proportion of them has been characterized of their involvement in cell wall components synthesis, catalytic processes, signaling transduction, stress and pathogen response, and so on (Zhou, 2019b). But to date, no particular GPI-AP has been reported to regulate integument development in plants.

*SKU5-Similar* (*SKU5/SKS*) gene family, named after *SKU5*, which has been reported to be involved in twisting bias of root growth in *Arabidopsis*, contains 19 members (Rutherford and Masson, 1996; Sedbrook et al., 2002). *SKS1, SKS2, SKS3* and *SKU5* (*GPI-SKU5/SKS*) within the same subgroup of this family could encode GPI-anchored proteins, and they were reported by us to be redundantly essential for root development through affecting anisotropic cell expansion in root and cell wall synthesis, and *SKS1* made significant contribution during this process (Zhou, 2019a; Zhou, 2019c).

In this study, we further investigated the involvement of this subgroup of *GPI-SKU5/SKS* genes in reproductive processes of *Arabidopsis. Loss-of-functions* redundantly and drastically reduced their seed production, which was demonstrated to be due to maternal-dependent ovule abortions. In those aborted ovules, the extension of their maternal integuments was disturbed and resulted in physical restriction to female gametophytes, which lead to their exposure to out space and consequent female gametophytic abortion. Although those less defective ovules could be fertilized and develop into seeds, their seed morphogenesis was largely affected as the aberrant integument/seed coat development.

The interference on integument development seemed due to similar defective polar expansion of integument cells as we described in their root cells previously, which is very surprising, as to our knowledge, none of gene has been reported to interfere polar expansion on both root and integument cells besides a series of MAPK components.

Our study firstly reported the crucial involvement of particular GPI-APs, GPI-SKU5/SKS proteins, in reproductive processes through regulating maternal integument development, which increased functional diversity of GPI-APs in *Arabidopsis*. Surprisingly, this group of GPI-APs could regulate cell polar expansion in both root and integument the same as a series of MAPK components.

## MATERIALS AND METHODS

### Pseudo-Schiff Propidium Iodide Staining

Ovules and seeds were harvested and then fixed in greenfix solution (Diapath) for 1 to 3 h at room temperature. Fixed samples were transferred into water and kept at 4°C for up to 7 d. The samples were rinsed once more with water and incubated in a 1%SDSand 0.2 N NaOH solution at 37°C for 1 h. The samples were rinsed in water and incubated in a a-amylase solution (0.4 mg/mL a-amylase, 20 mM NaPO4, and 6.7mMNaCl, pH 6.9) at 37°C for 2 h. The samples were rinsed in water and incubated in a 12.5% bleach solution (1.25% active Cl^−^) for 10 to 15 min. The samples were rinsed in water and then transferred to a 1% periodic acid solution at room temperature for 20 min. The samples were rinsed in water and incubated in a Schiff reagent and propidium iodide solution (100 mM sodium metabisulfite, 0.15 N HCl, and 100 mg/mL propidium iodide) until plants were visibly stained. The samples were rinsed in water and then incubated in achloral hydrate solution (4gchloral hydrate, 1mL glycerol, and 2 mL water) at room temperature for 30min (Fiume et al., 2017; Xu et al., 2016). Finally, the samples were transferred onto microscope slides for confocal laser scanning microscopy analysis (Leica SP5)

### Mature embryo sac observation

The oldest closed flower buds of a given inflorescence was emasculated and harvested two days later. The collected pistils were fixed in ethanol: acetic acid (9:1) solution overnight, and followed by two washes in 90% and 70% ethanol. Then they were cleared by a chloral hydrate:glycerol:water solution (8:1:2, w:v:v) shortly before observed under DMI6000 microscope system with white light (Gross-Hardt et al., 2007; Yadegari et al., 1994).

### Plant materials and growth conditions

*Arabidopsis.Thaliana Wassilewskija* (*Ws*) and columbia accession was acquired as wild type. Mutants acquired in the study were all in *Ws* background. *T-DNA* insertion lines, *sks1* (FLAG_521F09), *sks2* (FLAG_415F09), *sku5* (FLAG_386B03) were obtained from Arabidopsis Thaliana Resource Center of Institute Jean-Pierre Bourgin (IJPB) and *sks3* (SALK_067925). Seeds were surface sterilized and plated on 1/2MS medium containing 1% agar. After 48 hours vernalization at 4°C in dark, they were plated at 22°C under continuous light. About 7 days growth, seedlings were transferred into soil in green house. Primers for genotyping has been show in Table S1.

## RESULTS

### Maternally affected seed production in *gpi-sku5/sks* mutants

Characterization of various combinations of *null* mutations of members of the *GPI*-*SKU5/SKS* genes revealed their severe defects in seed production, and this group of genes seemed play redundant roles in this process (Fig.S1). Among *gpi*-*sku5/sks* triple and quadruple mutants, those with *SKS1* mutation generated shorter mature siliques (Fig.1A) containing a large proportion of small, wrinkled and white ovules that failed to develop into seeds (Fig.1B and 1C). Interestingly, together with quadruple mutant, *sks1;sks2;sks3* triple mutant was nearly infertile due to those failures of ovule development, and bare seed was produced.

**Fig1.**
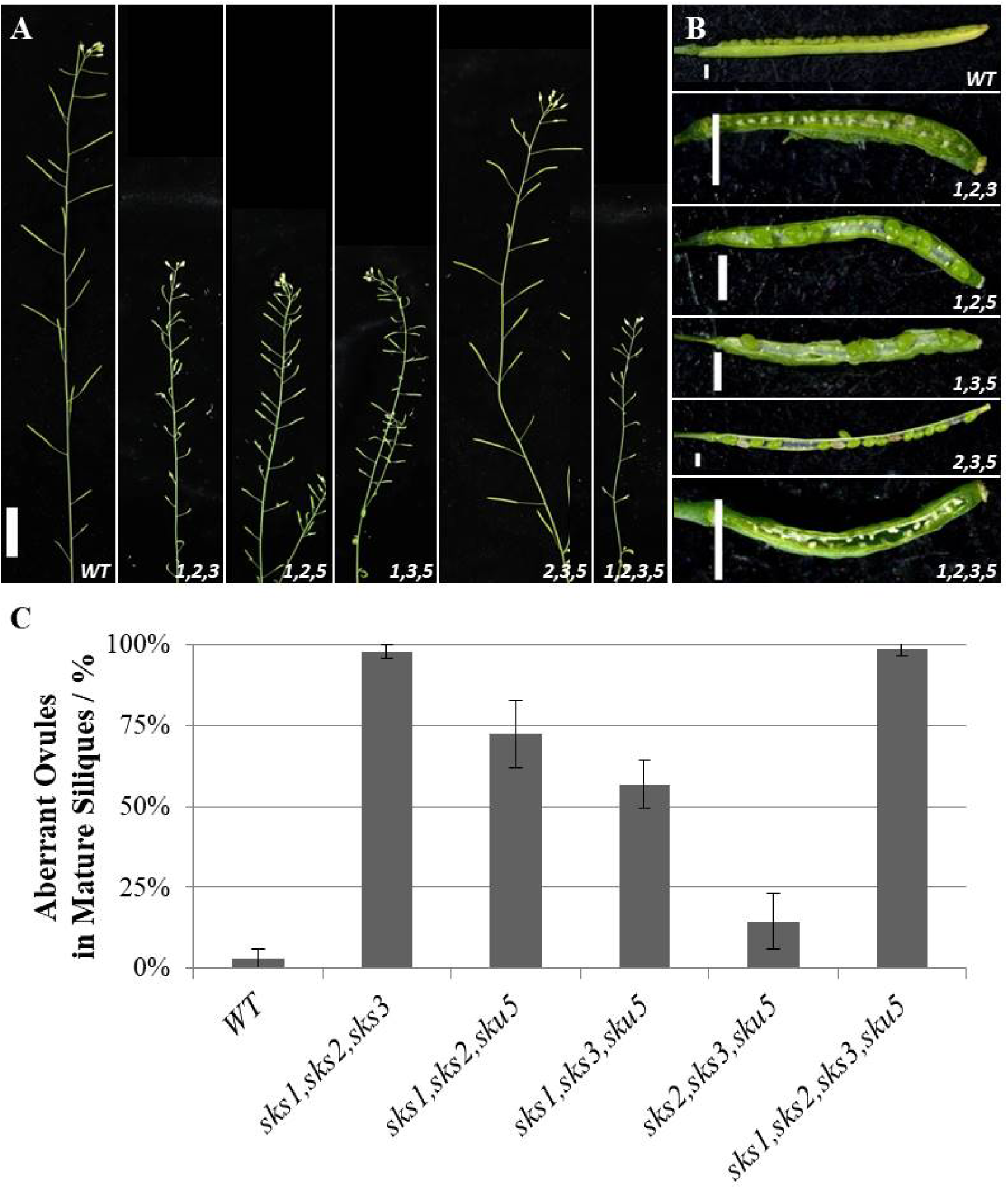
Dramatic decreased seed production in *gpi*-*sku5/sks* mutant. **(A)**, Mature siliques generated by wild-type and *gpi-sku5/sks* mutant lines. From Left to Right: Wild-type, *sks1,sks2,sks3*; *sks1,sks2,sku5*; *sks1,sks3,sku5*; *sks2,sks3,sku5* and *sks1,sks2,sks3,sku5*; bar, 2cm. **(B)**, Dissected siliques from *wild-type* and *gpi-sku5/sks* mutants. From Top to Bottom: wild-type, *sks1,sks2, sks3*; *sks1,sks2,sku5*; *sks1,sks3,sku5*; *sks2,sks3,sku5* and *sks1,sks2,sks3,sku5*; bar, 1mm. **(C)**, Proportion of white, small, and wrinkled undeveloped ovules among all ovules/seeds in mature green siliques from triple and quadruple *gpi-sku5/sks* mutants were shown. N=15 siliques, standard errors were indicated.

To determine the origin of such a phenotype, we analyzed the segregation of the quadruple mutant allele in the progeny of *sks1*+*/-;sks2;sks3;sku5* plants. A gametophytic- or zygotic-caused fertility due to *GPI-SKU5/SKS* mutations would result in rare detection of quadruple mutant seeds. By contrast, 20.69% (n=116) seedlings germinated from seeds generated by *sks1*+*/;sks2;sks3;sku5* plants were quadruple homozygous, which suggested the major contribution made by maternal tissue in this process.

According to our previous quantitative PCR (Zhou, 2019c) and *in silico* data (Fig.S2 and S3), unlike the specific expression of *SKS2* in reproductive organs, *SKS1, SKS3* and *SKU5* ubiquitously expressed at different level among the whole developmental progresses. Surprisingly, in spite of the significant contribution for reproduction, *SKS1* was expressed at relatively low level in reproductive organs.

Altogether, these data indicated that, *GPI*-*SKU5*/*SKS* genes played redundant and crucial roles in seed production, potentially through regulating ovule or seed development, and *SKS1* made significant contribution. By genetic analyses, defective maternal tissues were determined to be majorly responsible in this process.

### Drastic decreased seed production was due to ovule abortion in *gpi-sku5/sks* mutants

To further investigate the maternal origin of the *gpi*-*sku5/sks* mutant phenotype, the ovules from flowers just blooming were collected and observed using the modified pseudo-Schiff propidium iodide (mPS-PI) technique (Fiume et al., 2017; Xu et al., 2016). Surprisingly, more than 90% of the *sks1;sks2;sks3;sku5* ovules failed to develop an embryo sac (Fig.2B, and 2I), and the rest showed an exposed embryo sac that was not covered by the outer integument (Fig.2C and 2I), whereas both of them displayed irregular maternal integuments. The same defective ovules were also observed, at lower frequency, in *gpi-sku5/sks* triple mutant ovules (Fig.2D-2I) and the proportion of *sku5/sks* ovules that failed to form an embryo sac matched that of aborted ovules in mature siliques in their respective mutant backgrounds (Fig.1C and Fig. 2I).

**Fig2.**
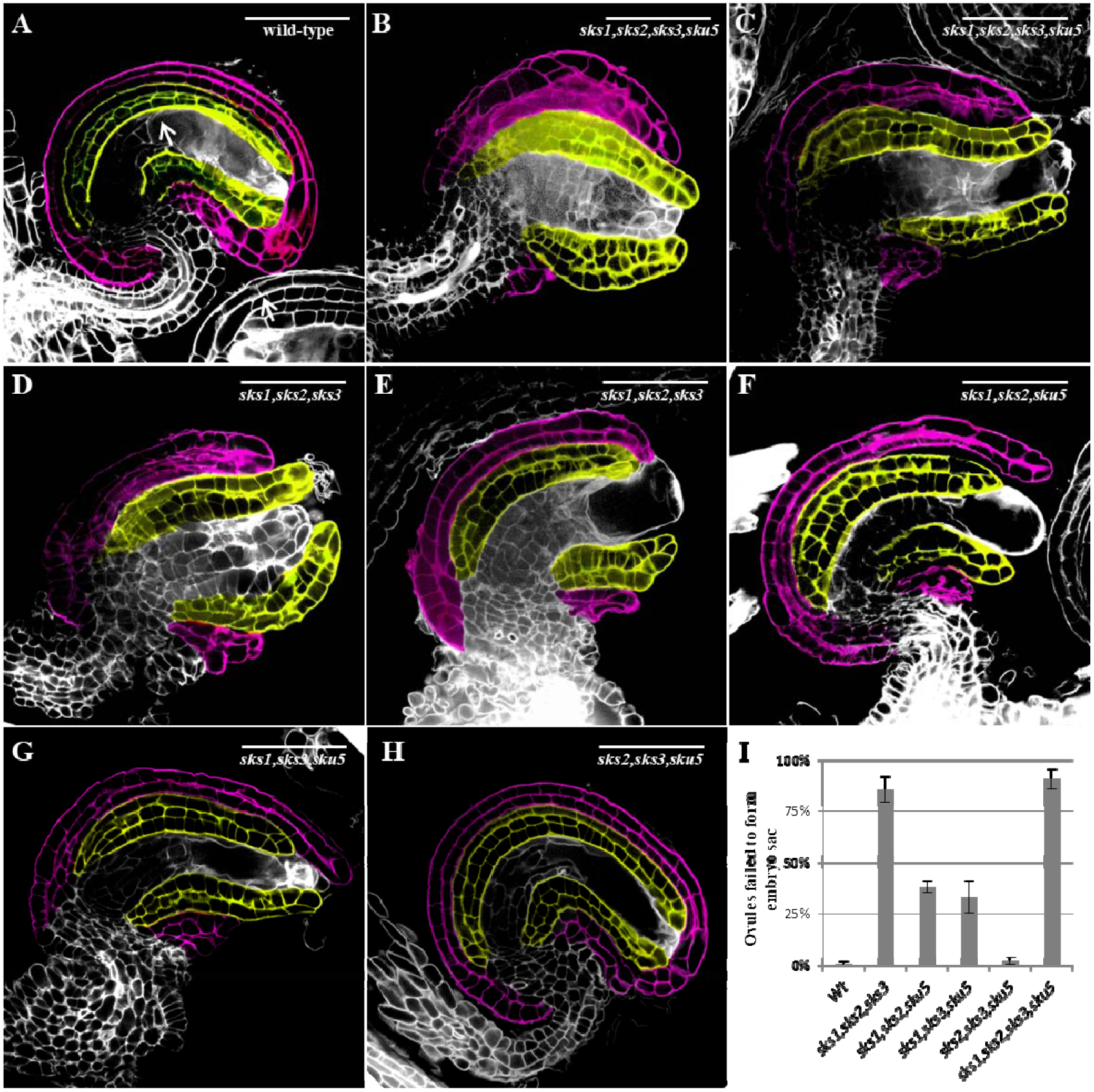
Defective ovules generated by *gpi-sku5/sks* mutants with false colors. Ovules were collected from just flowering siliques from (**A)** wild-type; (**B)** *sks1,sks2,sks3,sku5* that failed to form an embryo sac; (**C)** *sks1,sks2,sks3,sku5* that generated an aberrant embryo sac; **(D)** *sks1,sks2,sks3* that failed to form an embryo sac; **(E)** *sks1,sks2,sks3* that generated an aberrant embryo sac; **(F)** *sks1,sks2,sks5*; **(G)** *sks1,sks3,sks5*; and (**H)** *sks2,sks3,sks5*. To better visualized, inner integuments are colored in yellow, and outer integuments are colored in purple; bar: 50µM. **(I)** Proportion of ovules with embryo sac formation failures generated by various combinations of *sku5/sks* mutants. N=3 siliques, and the standard error bars were indicated.

It demonstrated that, the drastic decreased seed production observed in *gpi-sku5/sks* mutants maternally resulted from increased irregular ovules that failed to develop an embryo sac and the subsequent ovule abortion, and *GPI-SKU5/SKS* genes were redundantly involved in these processes.

### Female gametophytic abortion due to physical restriction resulted in ovule abortion in *gpi-sku5/sks* mutants

To follow the processes of their ovule abortion, the triple *null* mutants, *sks1;sks2;sku5*, which exhibited less defective ovules but be capable of generating reasonable seeds, were carefully studied. Our results showed a series of ovules and embryo sacs with various degrees of defections: slightly affected ovule of which integument could basically cover the embryo sac (Fig.3B); embryo sac were not well covered and partly exposed to external space (Fig.3C); embryo sac could barely covered by integument, but be exposed to external space as an outstretching bubble (Fig.3D); collapsing embryo sac at external space without protect by integument (Fig.3E); and ovule loss of embryo sac (Fig.3F).

**Fig3.**
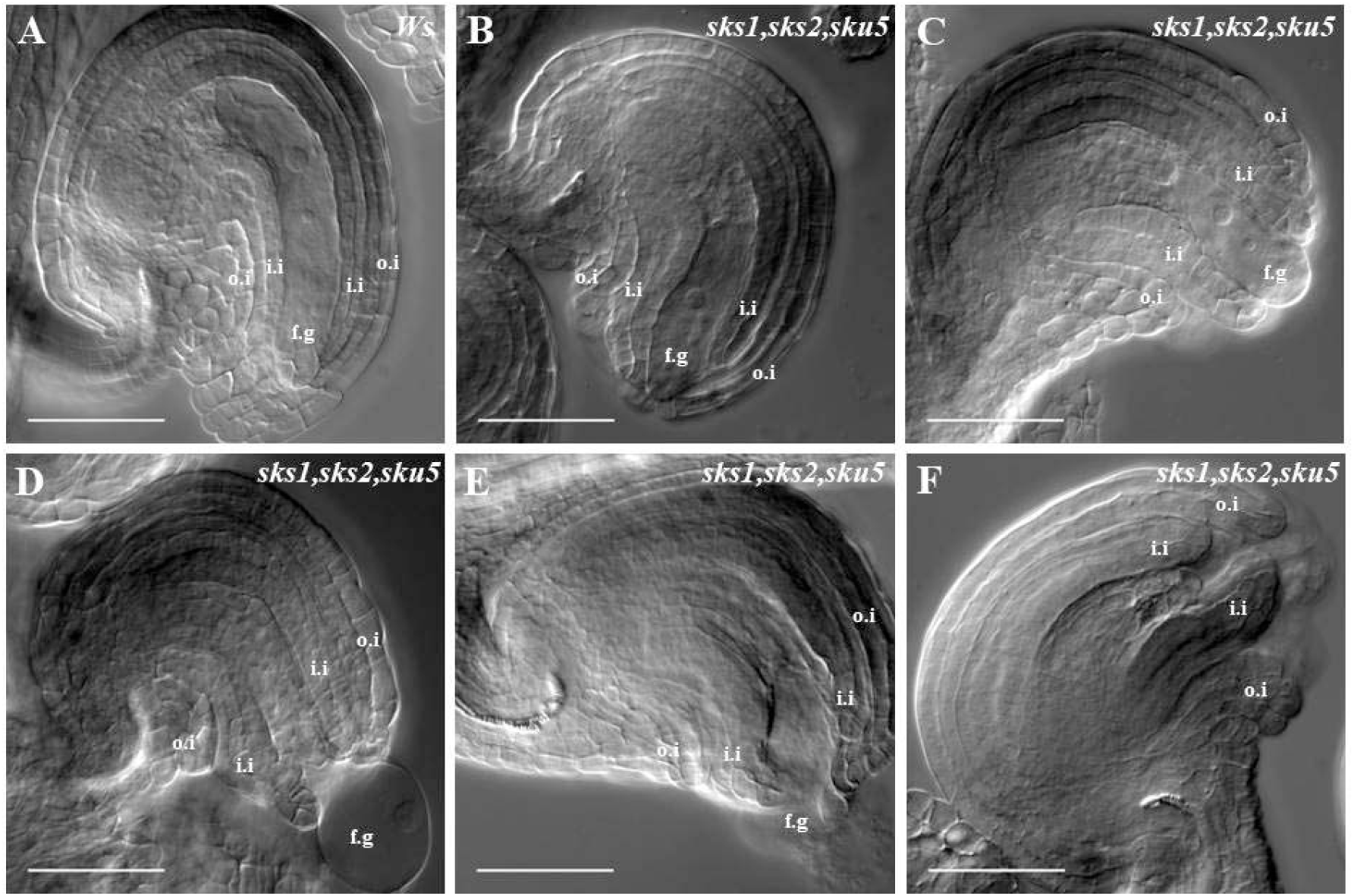
Female gametophytic abortion in *gpi-sku5/sks* triple null mutant line. Ovules with mature embryo sacs were collected and observed as described in “materials and methods”. **(A)**, wild-type; and a series of ovules and embryo sacs with various degrees of defections **(B-F)**; o.i: outer integuments, i.i: inner integuments, f.g: female gametophytes; bar, 50μm;

It indicated that, the female gametophytic abortion, which resulted from physical restriction due to undeveloped maternal integument, should be the main reason of ovule abortion in *gpi-sku5/sks* mutants.

### Fertilization, embryonic development and seed formation in *gpi-sku5/sks* mutants

Interestingly, those less affected ovules of *sks1;sks2;sku5* could be fertilized normally, however, due to defective integument development, inner integument could not be fully covered and normally curved by asymmetrically elongated outer integuments, which lead their embryos appeared mis-oriented (Fig.4A and 4B). Integuments develop into seed coats. Those mis-oriented embryos might develop at wrong position and break through the limit of defective seed coat and grow partially at external space (Fig.4C-4E). These defections certainly affected their seed morphogenesis largely, and exhibited irregularly shaped seed coat epidermal cells (Fig.4C-4E).

**Fig.4.**
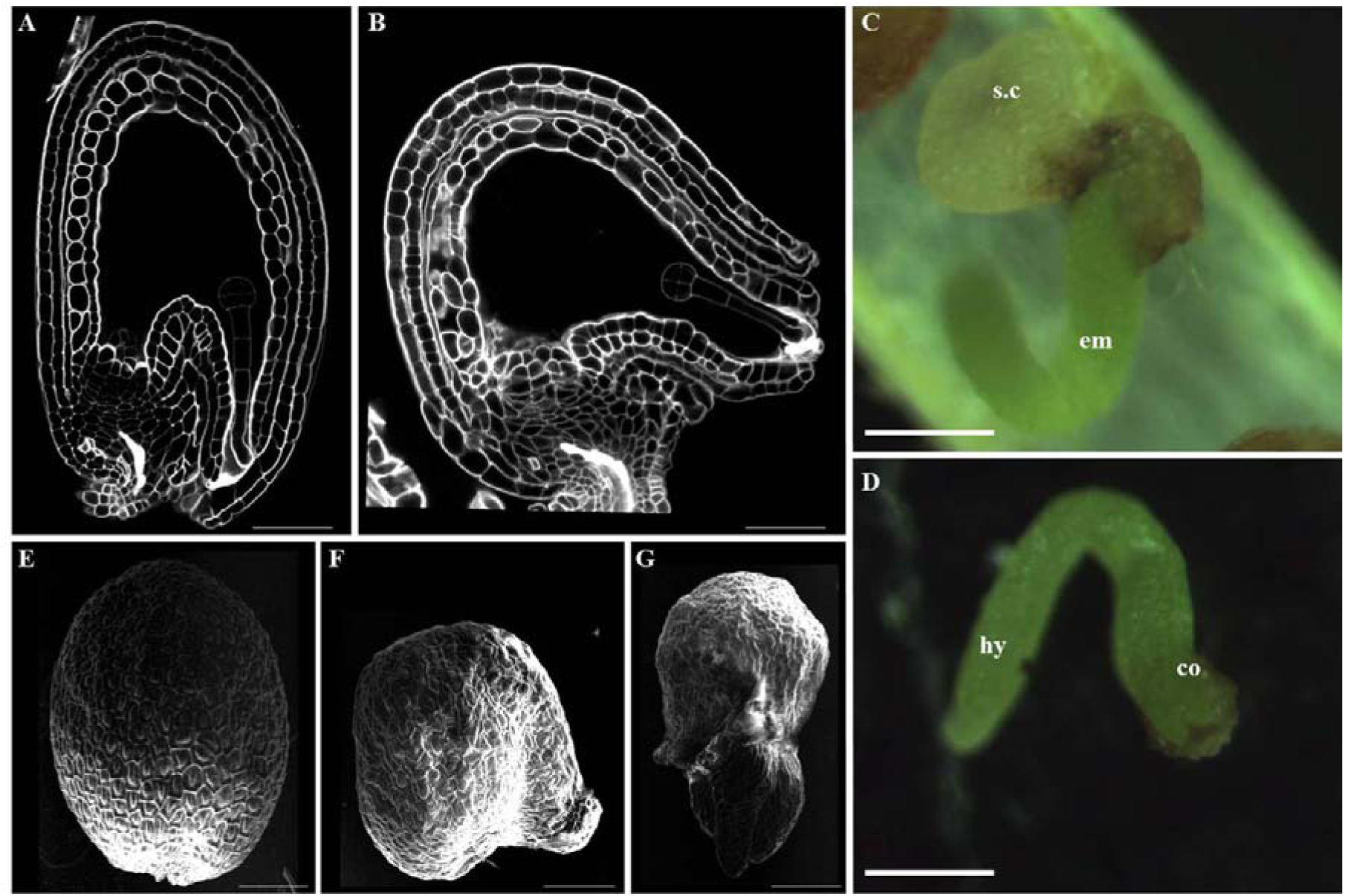
Embryonic development and seed formation in *gpi-sku5/sks* triple null mutant line. Embryos at pre-globular stage of **(A)** Wild-type and **(B)** *sks1;sks2;sku5* stained by mPS-PI technique and observed under confocal SP8; bar:50μm; **(C)** aberrant seed and **(D)** embryo separated from this seed generated by *sks1;sks2;sku5* mutant at mature embryo stage; s.c: seed coat; em: embryo; hy: hypocotyl; co: cotyledon; bar: 100μm; dry seeds from wild-type (**E**) and *sks1;sks2;sku5* mutant (**F** and **G**) under Scanning Electronic Microscopy; bar: 100μm.

It demonstrated that, the defections occurred on integuments of *gpi-sku5/sks* mutants also largely affect their seed formation and seed morphogenesis.

### Maternal regulation of seed morphogenesis in *gpi-sku5/sks* mutant lines

The same affected seed formation and morphogenesis were also found in other *gpi-sku5/sks* mutants that generated defective integuments (Fig.5A-5F), which are irregularly shaped, and some time with tissue protrusion (Fig.5H-5J). Interestingly, the affection on seed formation and morphogenesis was so strong that all seeds generated by mutants with aberrantly developed integuments were irregularly shaped (Fig.5K).

**Fig5.**
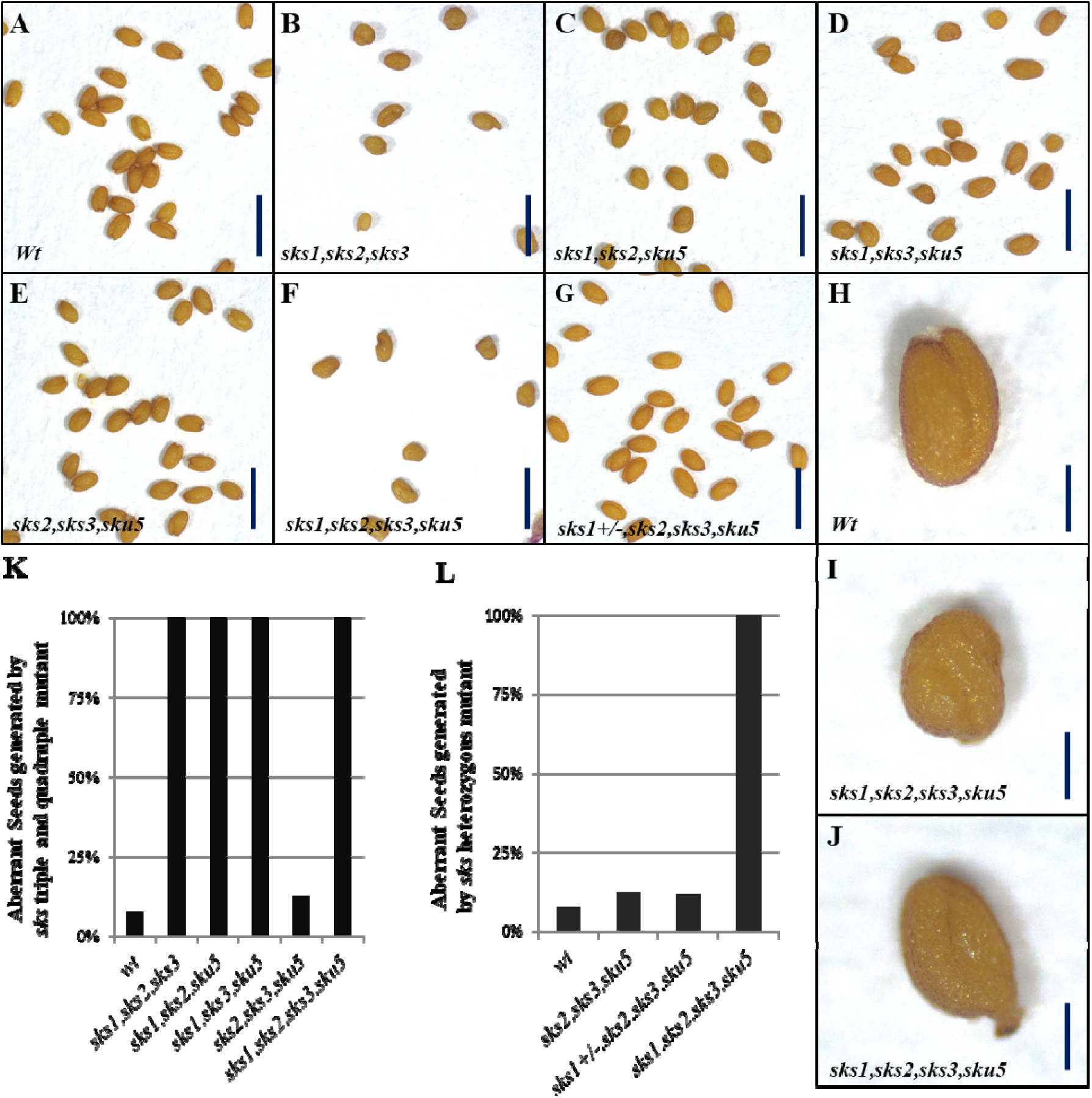
Aberrant seed morphogenesis in *gpi*-*sku5/sks* mutant and seeds segregation generated by *sks1*+*/-,sks2,sks3,sku5* heterozygotes. Mature seeds generated by **(A)**,wild-type **(B)**, *sks1,sks2,sks3* **(C)**, *sks1,sks2,sku5*; **(D)**, *sks2,sks3,sku5* **(E)**, *sks1,sks2,sks3,sku5* **(F)**, *sks1*+*/-,sks2,sks3,sku5* **(G)**, *sks1*+*/-,sks2,sks3,sku5*, bar: 1mm; typically seed morphogenesis of (**H**), wild-type and **(I-J)**, *sku5/sks* quadruple mutant, bar,0.2mm; **(K)**, proportion of aberrant seeds generated by all *sku5/sks* triple and quadruple mutants; **(L)**, proportion of aberrant seeds generated by triple, heterozygous and quadruple *sku5/sks* mutants. wild-type (n=375); *sks1,sks2,sks3*(n=11); *sks1,sks2,sku5* (n=234); *sks1,sks3,sku5* (n=322); *sks2,sks3,sku5* (n= 226), *sks1*+*/-,sks2,sks3,sku5* (n-347) and *sks1,sks2,sks3,sku5*(n=10).

To determine the contribution of maternal integument development regulated by *GPI-SKU5/SKS* genes in this process, the segregation of *sks1* mutant allele in the progeny of *sks1*+*/-;sks2;sks3;sku5* plants was characterized, which showed that, the aberrant seed formation and morphogenesis were only determined by the affected integuments, but not embryonic reason (Fig.5L).

These data indicated that, integument development regulated by *GPI-SKU5/SKS* genes were crucial during reproductive processes from gametophytic development to seed formation.

## DISCUSSION

### *GPI-SKU5/SKS* regulate maternal integument development in *Arabidopsis*

Integuments were totally maternal tissues and composed by special double-layer cylinder structures, inner- and outer-integument; after initiation on chalaza, the outer integument asymmetrically grew and eventually covered the inner integument and embryo sac, which determined the curvature of inner integument and shaped the ovule (Gallagher and Gasser, 2008; Gasser and Skinner, 2019; Kunieda et al., 2008; Villanueva et al., 1999). Integument development was believed regulated by a series of signaling pathways, including cell surface receptors (Jorda et al., 2016; Lee et al., 2015; Pillitteri et al., 2007; Terpstra et al., 2010), their intracellular signaling MAP Kinases (Bush and Krysan, 2007; Lopez-Bucio et al., 2014; Meng et al., 2012; Wang et al., 2008; Wang et al., 2007), and nucleus transcriptional factors (Elliott et al., 1996; Gaiser et al., 1995; Gallagher and Gasser, 2008; Gasser and Skinner, 2019; Kunieda et al., 2008; Nole-Wilson and Krizek, 2000; Villanueva et al., 1999). Although fertilization and seed production was largely affected when GPI moieties biosynthesis was interrupted in *Arabidopsis* (Bundy et al., 2016; Gillmor et al., 2005), the mechanism has not been revealed. Here, we firstly reported the involvement of particular GPI-anchored proteins in seed production and seed morphogenesis through regulating integument development.

### Defective integument generated by *gpi-sku5/sks* mutants

Through our genetic and microscopy analyses, both drastic decrease of seed production resulting from ovule abortion and aberrant seed morphogenesis resulting from defective seed coat formation found in *gpi-sku5/sks* mutants were determined to be due to irregularly developed integument. In those mutants, both inner and outer integument were affected, however, defective outer integument, which was supposed to cover the inner integuments, to enclose the embryo sac and to shape the ovules, contributed more in this process. It made inner integument growing symmetrically, and embryo sac exposing to external space, which resulted in loss of female gametophytic due to physical restriction and consequent ovule abortion. Interestingly, those less affected ovules could be fertilized and develop into seeds, however, mis-localization of embryos, or insufficient seed coat that both due to defective integuments would result in post-embryonic abortion or aberrant seed morphogenesis. Here, our study clearly demonstrated the crucial roles *GPI-SKU5/SKS* genes played during integument development, and how they affected seed production and morphogenesis in *Arabidopsis*.

### Disturbed cell polar expansion in *gpi-sku5/sks* mutants

In *gpi-sku5/sks* mutants, defective integuments seemed due to their disturbed cell polar expansion (Coen and Magnani, 2018; Endress, 2011; Enugutti et al., 2012; Enugutti and Schneitz, 2013; Gaiser et al., 1995; Scholz et al., 2019), where cells from both inner and outer integument seemed shorter, thicker, and aberrantly shaped. It largely disturbed their elongation and extension, where *GPI-SKU5/SKS* played redundant roles and *SKS1* made significant contribution. The same redundancy and significant contribution of *SKS1* in root cell polar expansion were reported by us recently, which exhibited similar shorter, thicker and aberrantly shaped cells (Zhou, 2019c).

But surprisingly, to our knowledge, none of gene has been reported to interfere polar expansion on both root and integument cells besides a series of MAPK components, YODA(MAPKKK)-MEK4/5(MAPKK)-MPK3/6(MAPK) cascade. And interruption of this cascade resulted in similar defective polar expansion in both roots and integuments (Bush and Krysan, 2007; Lopez-Bucio et al., 2014; Smekalova et al., 2014; Wang et al., 2008; Wang et al., 2007; Zhang et al., 2017).

### The potential involvement of GPI-SKU5/SKS proteins in signaling pathway

As been reviewed recently, GPI-Anchored Proteins in *Arabidopsis* broadly participate in various biological processes, including signaling transduction through associating with cell surface receptors and/or extracellular ligands, to mediate developmental processes, stress and immune response, and so on (Zhou, 2019b). Interestingly, SKU5 was reported to directly interact with the synthesized functional Carboxyl-terminal region of Auxin Binding Protein 1 (Shimomura, 2006), that was recently believed to be a potential extracellular ligand that could bind Auxin and mediate extracellular auxin signaling through binding auxin and being recognized by cell surface reporter TMK1, then activate quick auxin response (Cao et al., 2019; Xu et al., 2014). Additionally, exogenous auxin could rapidly activate YODA(MAPKKK)-MEK4/5(MAPKK)-MPK3/6(MAPK) cascade (Mockaitis and Howell, 2000; Pillitteri et al., 2007; Smekalova et al., 2014), of which interruptions were reported to exhibit similar defective roots and integuments as *gpi-sku5/sks* mutants.

Altogether, GPI-SKU5/SKS associate with Auxin-related extracellular polypeptide/ligand ABP1, which could be recognized by cell surface receptor kinase TMK1 and activate intracellular signaling components; and their *loss-of-functions* exhibit similar defections as interrupted intracellular MAPK cascade components. It exhibits us a hypothesis that, GPI-SKU5/SKS may work as typical GPI-APs in signaling transduction during regulation of cell polar expansion in both root and integument cells.

## Supplemental Materials

**Fig.S1.**
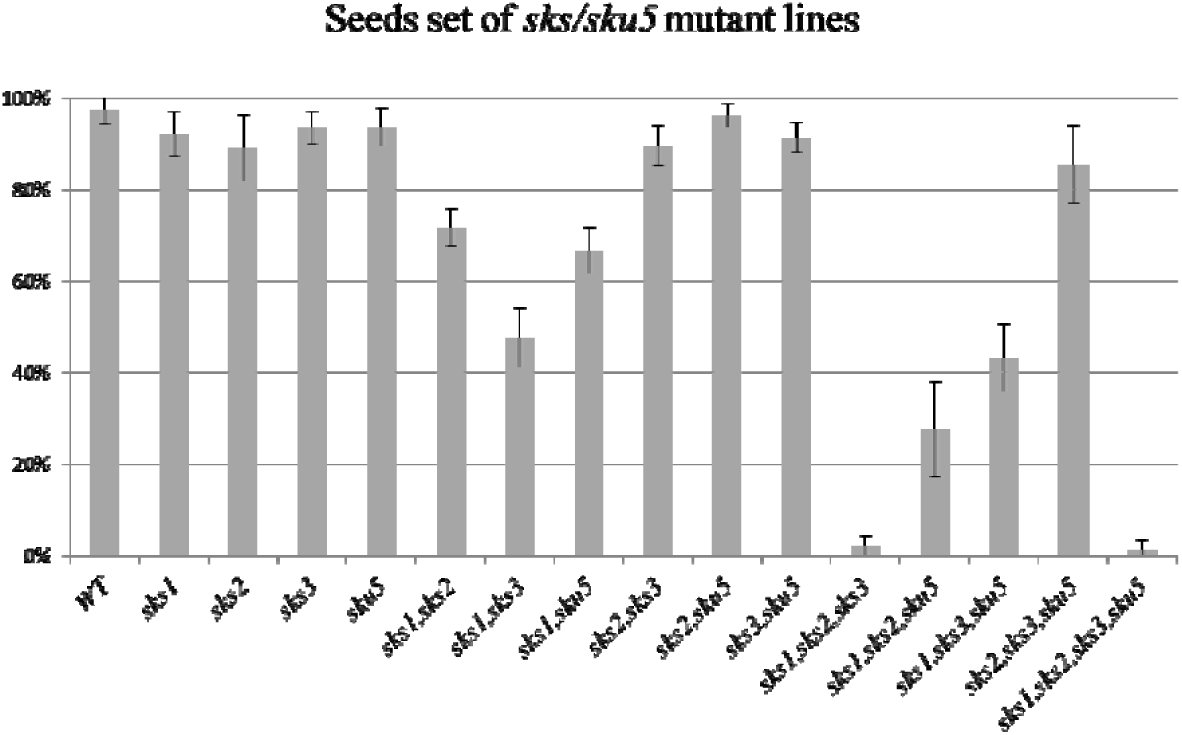
Seed sets of all *gpi-sku5/sks* mutants. Proportion of mature seeds among all ovules/seeds in mature green siliques from wild-type and *gpi-sku5/sks* mutants were shown. N=15 siliques, and standard errors were indicated.

**Fig.S2.**
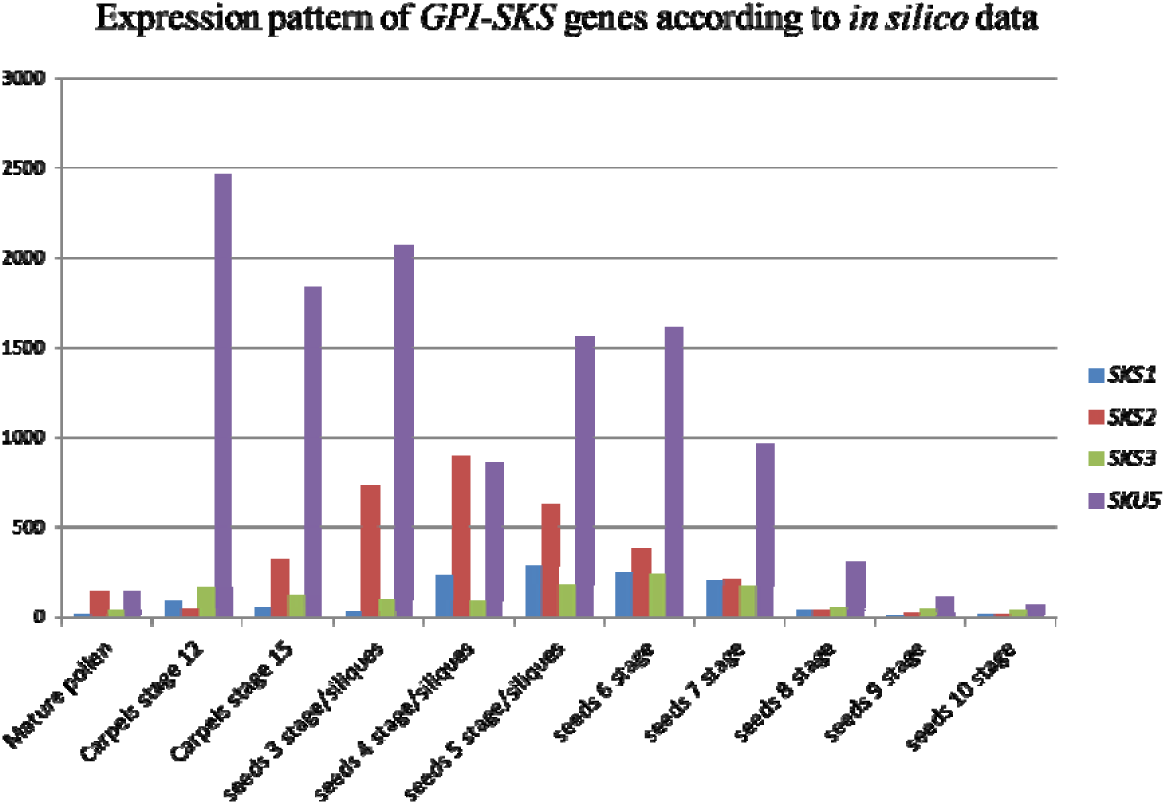
Expression level of *GPI-SKU5/SKS* genes in reproductive processes of *Arabidopsis.* The expression patterns of *GPI-SKU5/SKS* genes in reproductive organs based on microarray data from AtGenExpress Visualization Tool (AVT), were shown. Different genes were show with different colors, different developmental stages were shown at X-axis and the relative expression levels were shown at Y-axis.

**Fig.S3.**
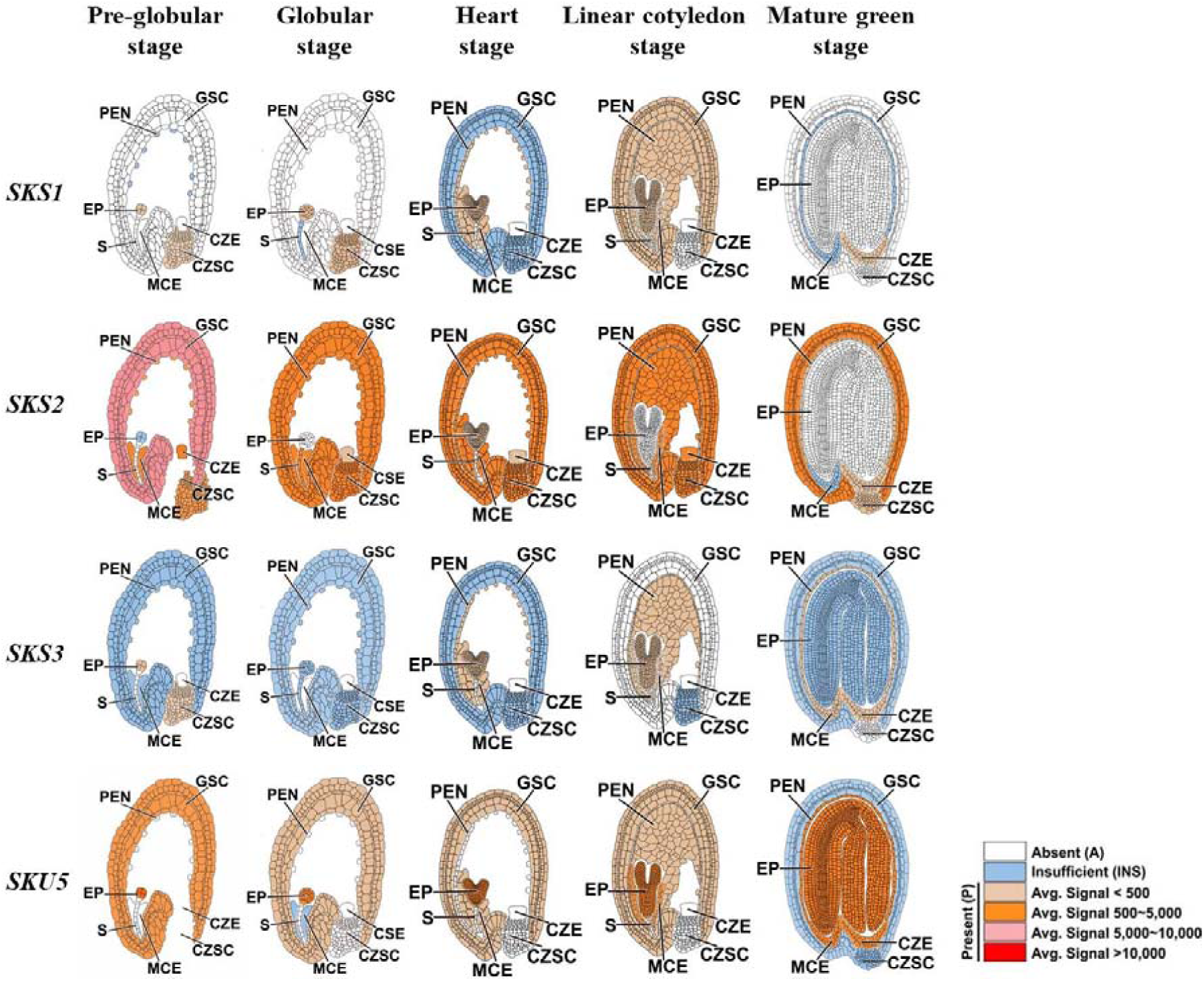
Exprssion level of *GPI-SKU5/SKS* genes during embryo development of *Arabidopsis*. Expression pattern of *GPI-SKS* genes during different embryo developmental stages based on microarray data from database http://seedgenenetwork.net/ were shown in different colors, which represented their expression levels. PEN, Peripheral Endosperm; GSC, General Seed Coat; CZE, Chalazal Endosperm; CZEC, Chalazal Seed Coat; MCE, Micropylar Endosperm; S, Suspendsor; EP, Embryo Proper;

**Table.S1.**
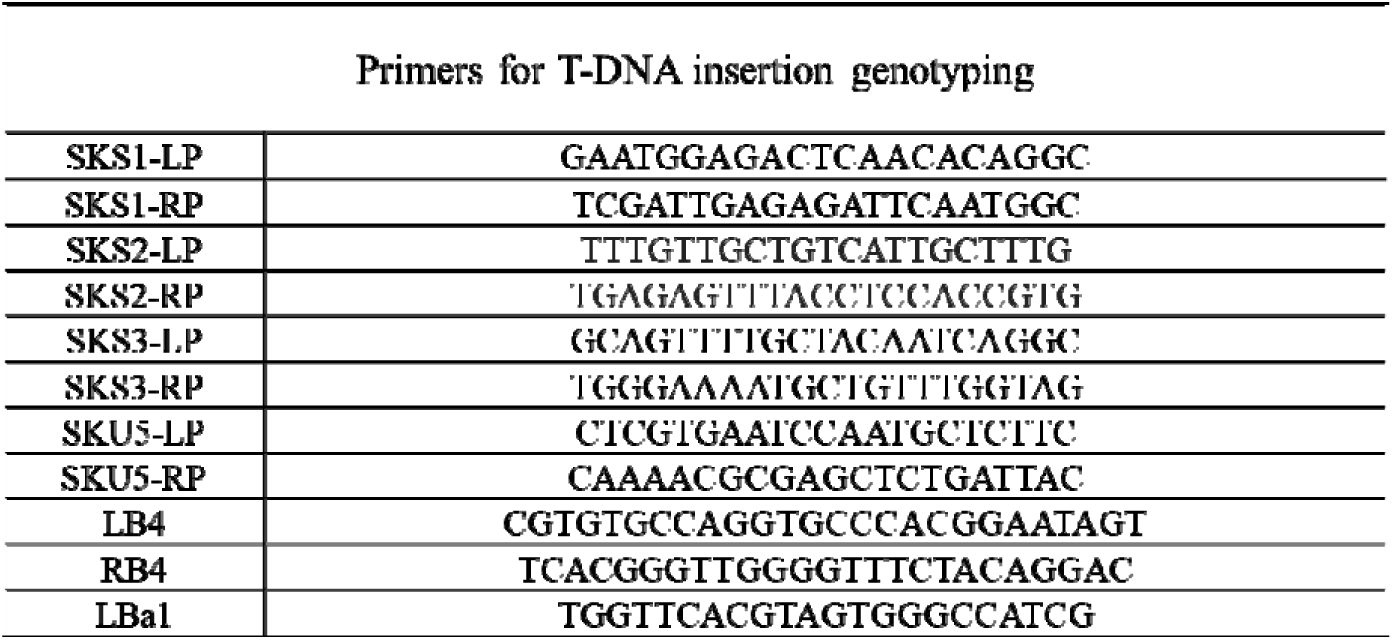
Primers being utilized for genotyping T-DNA insertions in *gpi-sku5/sks* mutants were shown.

